# Machine Learning Enables Viral Genome-Agnostic Classification of RNA Virus Infections from Host Transcriptomes

**DOI:** 10.1101/2025.09.11.675678

**Authors:** Benjamin Kaza, Amandine Gamble, Stephanie N. Seifert, Timothée Poisot

**Affiliations:** Department of Public and Ecosystem Health, Cornell University, Ithaca, NY, USA; Department de Sciences Biologiques, Université de Montréal, Montréal, QC, Canada; Paul G. Allen School for Global Health, Washington State University, Pullman, WA, USA

## Abstract

Targeted PCR diagnosis of RNA viruses is sequence dependent, meaning that the accuracy of the assay depends on the identity of the viral sequence. However, sequence-targeted assays can miss novel or divergent viruses. We test whether the host transcriptome alone can classify RNA virus infections without using viral sequences. Using publicly available data on Huh7 and Calu-3 cells experimentally infected with diverse negative-sense RNA, we evaluate two host-derived feature sets (differential expression dataset of identified genes and alignment-free nucleotide k-mer spectra) and compare hierarchical clustering with Random Forests. Across datasets and timepoints (12 to 24 hpi), Random forests accurately distinguished cells infected with different viruses. Controls with label permutation and read shuffling across experimental conditions established non-random performance. Influenza A infections exhibited the strongest, most distinct signatures, whereas Ebola and Lassa virus responses were subtler yet still classifiable. These results show that host-only transcriptomics encodes virus-specific, complex signals that machine learning can exploit, enabling genome-agnostic classification of infection statuses. This approach could aid early outbreak triage when primer based detection methods fail or viral genomes are unknown and complements sequence-based discovery.

## INTRODUCTION

Zoonotic viruses—those that spill over from animal reservoirs into human populations—pose a growing threat to global health. Anthropogenic global change and biodiversity loss are accelerating the frequency of spillover events by altering ecosystems, disrupting host-pathogen dynamics, and increasing contact between humans and wildlife^1,2^. While only a subset of animal viruses known to be are capable of infecting humans, those that do are disproportionately likely to cause severe disease and, in some cases, global pandemics^3–7^. Despite this risk, the vast majority of viruses circulating in wildlife remain undescribed^8–10^. Most have never been isolated, sequenced, or even detected^9–11^. This knowledge gap severely limits our ability to assess which viruses might pose a threat to public health. In addition, once in humans, many viruses may cause subclinical or misdiagnosed diseases (such as undiagnosed febrile diseases that could resemble relatively common diseases like malaria or influenza), further complicating efforts to identify them before they spread^12,13^.

Targeted PCR-based assays face similar challenges. These methods depend on primers designed to match conserved regions of known viral genomes. If a virus’s genome diverges too much from the expected sequence, primer binding may be inefficient or fail entirely, resulting in false negatives^14–17^. Even assays designed with degenerate primers—such as pan-family PCRs—can fail to detect divergent strains. For example, a second genotype of Hendra virus (HeV-g2) circulated undetected in a surveilled population of bats due to mismatches with primers designed for the original Hendra virus genome^18^. Similarly, a recombinant coronavirus (CCoV-HuPn-2018) that caused febrile illness in children in Malaysia and Haiti escaped detection by coronavirus-specific RT-PCR^19,20^. This is particularly concerning because Henipaviruses and Coronaviruses are both considered WHO priority pathogens that are flagged for their potential to cause pandemics with high fatality rates, and in these two (and potentially many more) cases they were missed^21^.

Modern virus discovery relies heavily on sequence-based methods (such as genome amplification by targeted PCR for detection and sequencing), particularly high-throughput sequencing and metagenomics^22,23^. These approaches enable the recovery of viral genome fragments directly from clinical or environmental samples without prior knowledge of the virus’s identity^24,25^. However, sequence-based discovery typically requires some level of similarity between the unknown virus and existing sequences in reference databases^26–28^. Highly divergent viruses may go undetected if they lack conserved genes or recognizable sequence motifs^29,30^. This is a particularly relevant problem for RNA viruses, which tend to have high mutation rates, genetic variability, and extreme expression differences between genes making it difficult to pick a good PCR target across families—and even within genera^31^. For example, transcription in Ebola virus (EBOV) follows a 3’ – 5’ gradient with the RNA-dependent RNA polymerase being the least expressed gene^32^. As such, there is an urgent need for virus detection strategies that do not depend on prior sequence information.

One alternative is to detect infection by measuring changes in the host cell, rather than detecting the virus directly. Viral infection can induce characteristic transcriptional responses in host cells^32,33^, which may serve as indirect markers of infection. Host-based diagnostic methods are already used clinically to distinguish between viral and bacterial infections by analyzing expression patterns of 29 genes that are up- or downregulated during infection^34,35^. Some studies have used short k-mers from viral genomes to train machine learning models that can classify viral infections from total RNA-seq data, without relying on alignment to reference genomes to determine the presence or absence of one type of viral infection or to discriminate between bacterial or viral infection^36–39^.

However, few studies have systematically tested whether host transcriptomic responses alone— without any viral sequence—can be used to classify infections across different viral families. It remains unclear whether the host response provides enough resolution to infer the identity or family of the infecting virus, especially across diverse cell types and timepoints.

In this study, we test the hypothesis that viral species can be inferred from host transcriptomic responses alone. We evaluate two feature types (short nucleotide k-mer spectra from total RNA and differential gene expression) and assess their utility for classifying viral infections using machine learning and hierarchical clustering. We focus on negative-sense RNA viruses infecting mammalian cell lines, using publicly available transcriptomics datasets^40,41^. Our panel includes experimental infections of Huh7 cells (a human hepatocyte immortalized cell line) and Calu-3 cells (a human lung epithelial immortalized cell line) with viruses including Influenza H1N1, H5N1, H7N7, H7N9, EBOV, Marburg virus (MARV), Respiratory Syncytial Virus (RSV), Nipah virus (NiV), Lassa virus (LASV), Rift Valley Fever virus (RVFV), and Sandfly Fever Sardinia Virus (SFSV). Across both cell types, we show that different viruses elicit distinct, classifiable host responses—supporting the possibility of sequence-independent viral classification using host transcriptional data alone.

## RESULTS

### Gene expression patterns elicited by infection vary in magnitude and differentiability

To test the hypothesis that each viral infection results in a differentiable response compared to other viruses, we performed hierarchical clustering of differential gene expression profiles from experimentally infected Huh7 cells at 12 and 24 hours post infection. In short, hierarchical clustering is a method for grouping similar data points into clusters by progressively merging the closest pairs to form a tree-like structure called a dendrogram. We used the Euclidean distance, which is a measure of the straight-line distance between two points in space, calculated as the square root of the sum of the squared differences between their corresponding coordinates^42^.

The magnitude and specificity of the host transcriptional response varied across viral infections, resulting in distinct patterns of gene expression. Hierarchical clustering using the top 100 differentially expressed genes effectively separated infected from uninfected samples (Figure 1). While some viral infections elicited sharply divergent transcriptomic profiles, others were more subtle and clustered closer to uninfected controls. Notably, infections with Influenza A viruses (H1N1 and H5N1) induced the most distinct expression patterns, clustering farthest from the uninfected samples and from other viruses.

**Figure 1:**
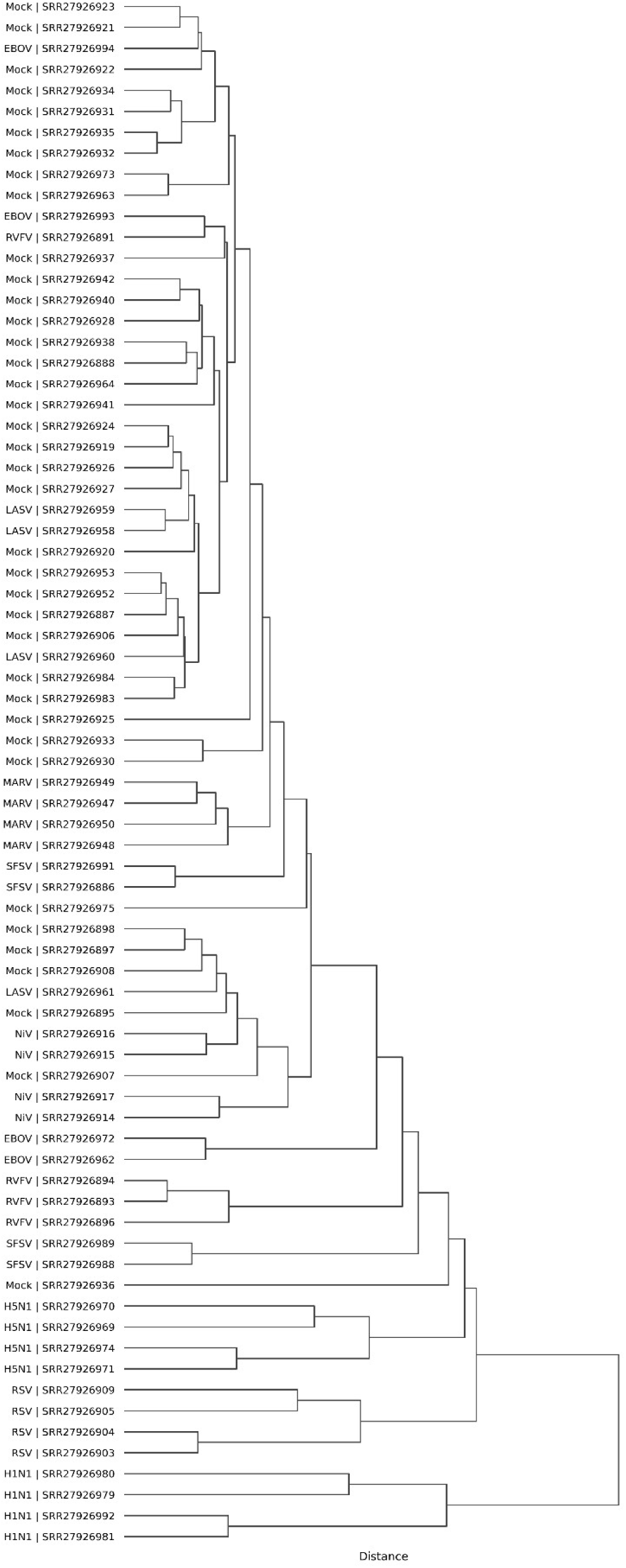
Hierarchical clustering of gene expression profiles from Huh7 cells 12 hours post infection

Several other viruses—including NiV, RSV, SFSV, RVFV, and MARV—also produced expression profiles distinguishable from uninfected controls, as their transcriptomes were placed outside of the subtree containing most mock-infected samples. In contrast, infections with EBOV and LASV produced host responses that were less distinguishable, with infected samples clustering within or adjacent to the subtree of uninfected controls (Figure 1).

The genes driving these distinctions were primarily involved in translation, immune response, and chromatin remodeling, followed by genes associated transport, cell signaling, and metabolism (Figure 2).

**Figure 2:**
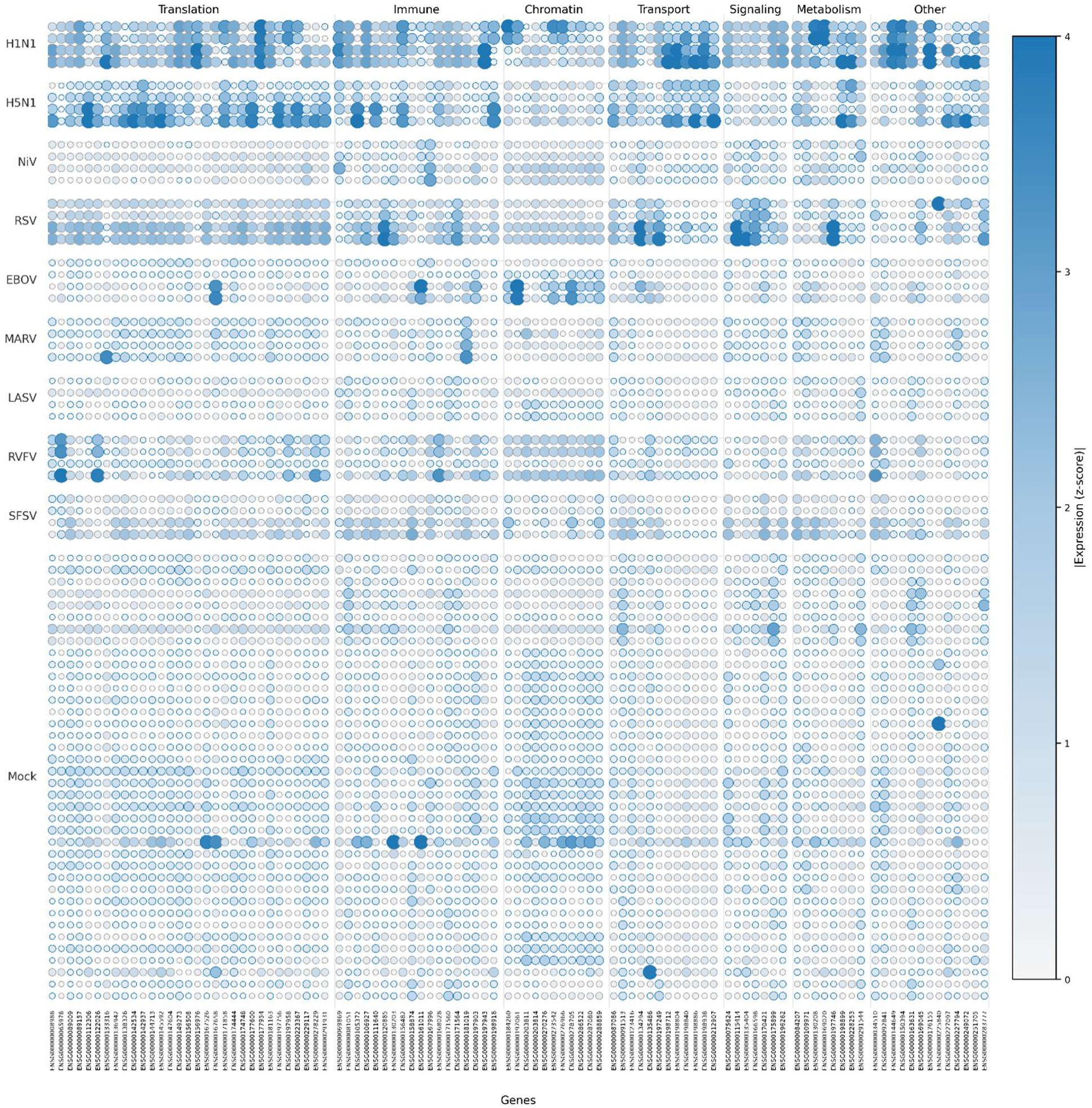
Z scores of top 100 differentially expressed genes in Huh7 cells infected with various negative-sense RNA viruses after 12 hours post infection and mock

A consistent feature across most infections was the downregulation of genes involved in translation, a known host antiviral response^43–45^; however, this trend was reversed in H5N1 and H1N1, and to a lesser extent RVFV, where translation-associated genes were upregulated relative to uninfected samples. Furthermore, only H5N1, H1N1, and MARV exhibited clustering patterns so distinct that their replicates grouped into clearly defined, virus-specific subtrees.

### The patterns of gene expression elicited by closely related Influenza A viruses are specific and differentiable by hierarchical clustering and random forest classification

To test whether Influenza A viruses consistently induced differentiable responses in a different cell type and across viral genotypes, we used experimental infection micro-array gene expression data from Calu-3 cells experimentally infected with H1N1, H5N1, H7N7, and H7N9 viruses^41^. We found that all of the four viruses were able to be classified by both Hierarchical clustering and random forest classification at 12 and 24 hours post infection (Figure 3).

**Figure 3:**
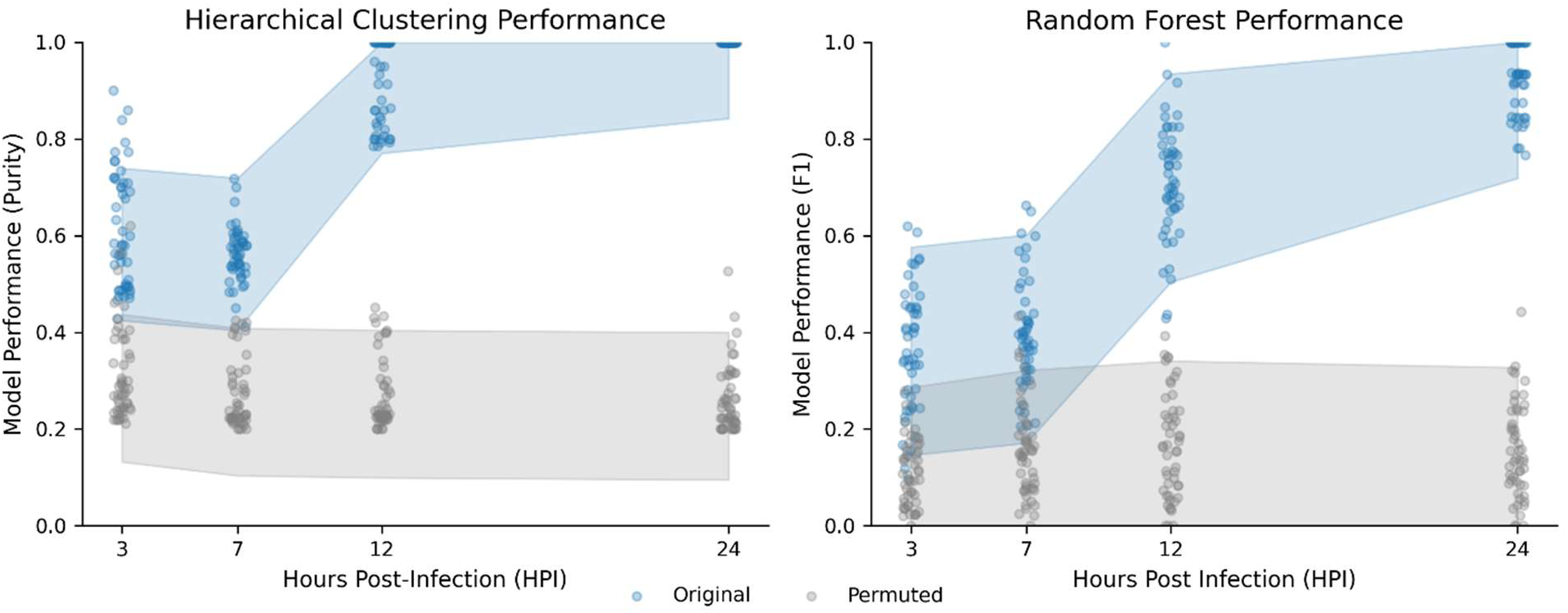
Hierarchical clustering Tree Purity and Random forest Classification F1 score of Differential Gene Expression Profiles originating from Calu-3 cells infected with various Influenza A viruses or mock at 3, 7, 12, and 24 hours post infection

### The distribution and composition of nucleotides in an infected cell varies in a virus-specific manner

To test whether the nucleotide composition of an infected cell can be used to predict which virus was infecting the cell, we compared experimental data to simulated parallel datasets where the nucleotides were randomly shuffled within each sequencing read or a dataset where the infection status label was randomly assigned to each kmer abundance profile.

Infection status was differentiable for kmer lengths 8 to 12 above random chance across all of the viruses tested via hierarchical clustering using Euclidean distance. Virally infected transcriptomes were differentiable to random forest classification above randomly assigned data for the entire k-mer range, and was differentiable to the model in contrast with nucleotide-shuffled data when the k-mer length was greater than 4 quantified by purity and F1 score which measure the accuracy of hierarchical clustering or random forests to group samples of the same category together, respectively (Figure 4).

**Figure 4:**
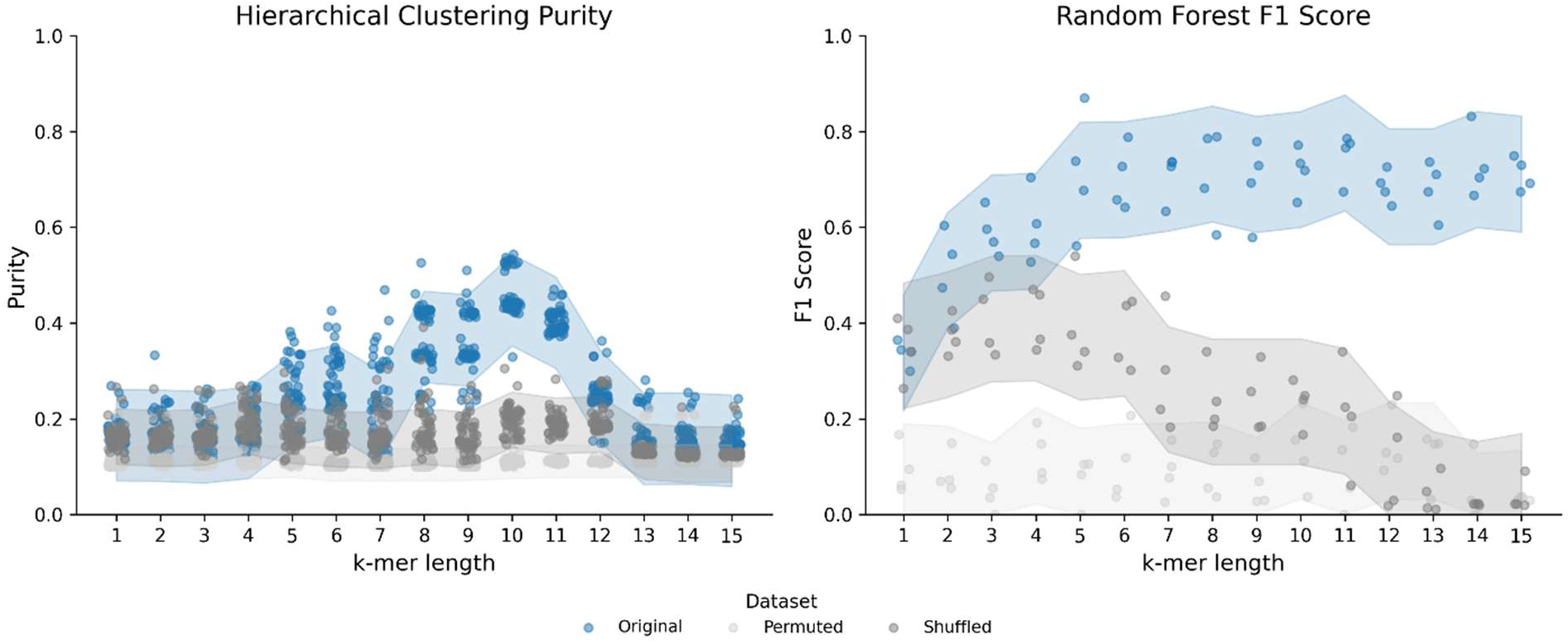
Performance of Hierarchical clustering measured by Tree Purity (a), and Random forest F1 Score (b), Recall (c), and Precision (d) on infected and uninfected Huh7 transcriptomes after 12 hours post infection and synthetic datasets where all of the reads in the original transcriptomes were randomly shuffled, or where each k-mer abundance profile was randomly assigned a virus (ie. permuted)

### The contribution of nucleotides from negative-sense RNA viruses drives the differentiability of k-mer abundance profiles in hierarchical clustering but not random forest classification

To test if the viral RNA itself drives the differentiability of the transcriptomes in Random forest classification and Hierarchical Cluster, we created a synthetic dataset where the viral reads were depleted.

In transcriptomes where reads originating from each virus were depleted, at no point in the kmer range were the transcriptomes differentiable above the shuffled or permuted data. In Random forest classification, kmers above length 2 were able to inform the model above random chance, but not to consistently high accuracy (Figure 5).

**Figure 5:**
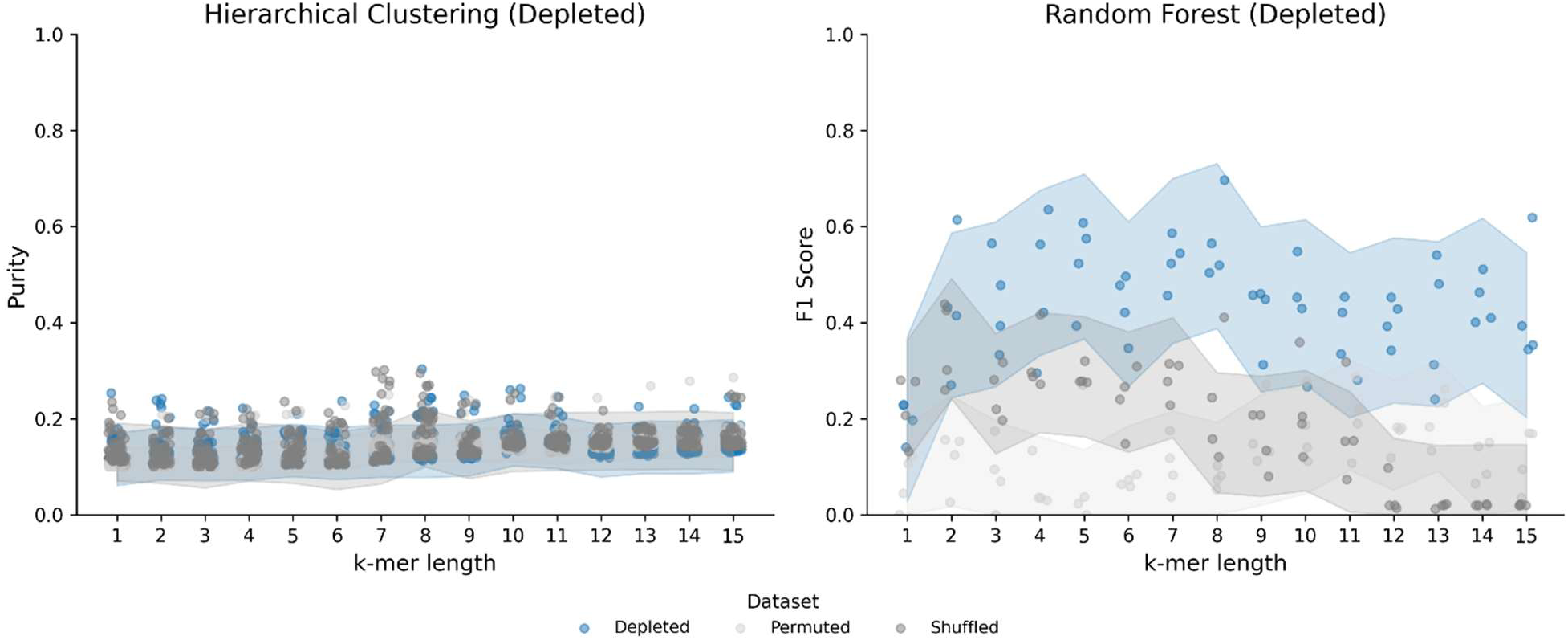
Performance of hierarchical clustering measured by tree purity (a) and random forest F1 score (b) on infected and uninfected Huh7 transcriptomes at 12 and 24 hours post infection that were computationally depleted of viral reads

### Enrichment of viral RNA in transcriptomic data enhances the differentiability to random forest classification and hierarchical clustering

When viral reads were doubled in the dataset, the resulting differentiability by hierarchical clustering and random forest classification increased. For hierarchical clustering, the data was differentiable above random chance over the k-mer range 4 – 12 and was differentiable by random forest classification for all k-mer lengths tested (Figure 6).

**Figure 6:**
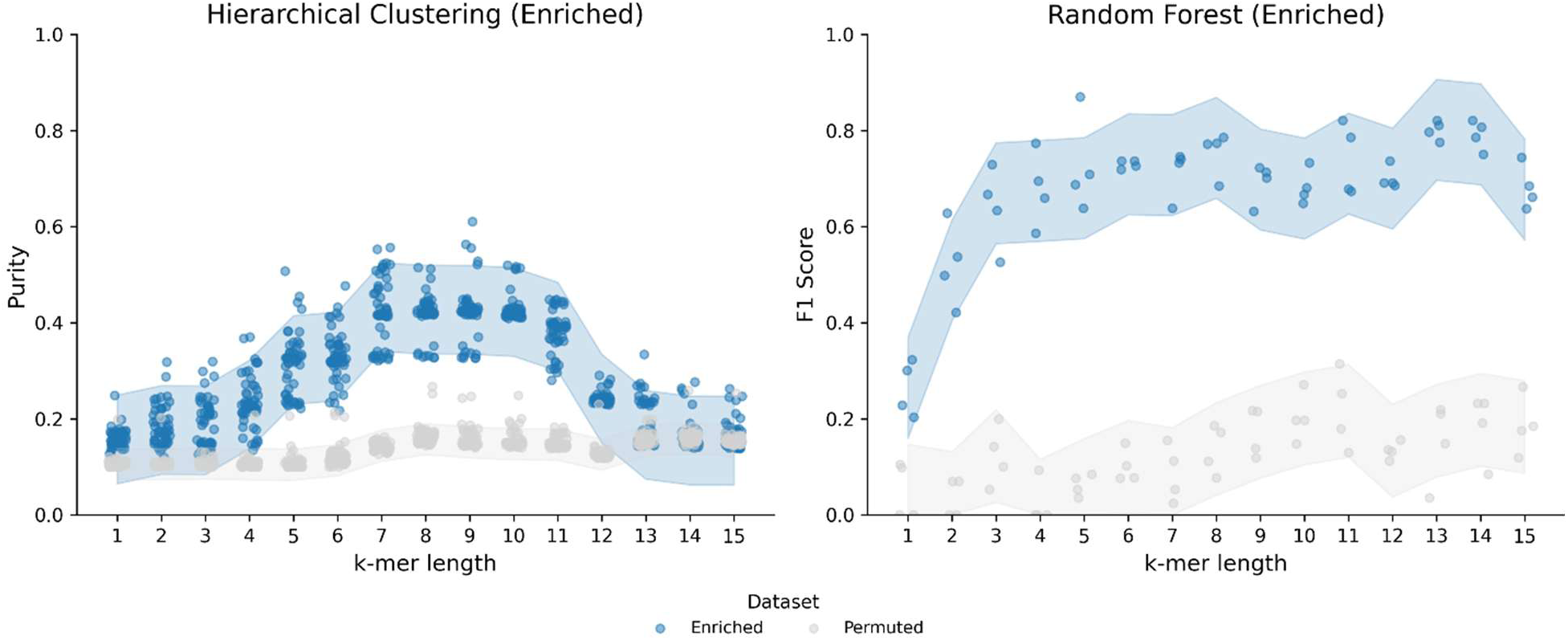
Performance of Hierarchical clustering measured by Tree Purity (a) and Random forest F1 Score (b) on infected and uninfected Huh7 transcriptomes at 12 and 24 hours post infection that were computationally enriched for viral reads

## DISCUSSION

To replicate and spread, all viruses must manipulate host cell biology^46^. At the cellular level, this often involves subverting host gene expression and regulatory networks to promote viral replication, suppress immune responses, and evade detection^46^. Viruses systemically manipulate the host’s immune system, inhibit antiviral signaling pathways, rewire host metabolism, or disrupt intercellular communication to maximize their replication fitness^47–49^.

Despite their compact genomes, RNA viruses encode multifunctional proteins that carry out both essential replication tasks and host antagonism^46^. These dual roles are a product of strong selective pressures: with limited coding space, many RNA viruses have evolved proteins and genome structures that both facilitate replication and interfere with host defenses. Viral replication itself often hijacks or suppresses core host processes at every stage—transcription, translation, signaling, and transport^47,50–52^.

In turn, host cells have evolved sophisticated and dynamic genetic mechanisms to detect and restrict viral infection. Mammalian cells encode hundreds of innate immune genes that are activated in response to viral invasion, though the specific pathways engaged can vary depending on the virus and host cell type^33,50–53^. Effective immune responses must be both rapid and specific: activating the appropriate antiviral programs without triggering unnecessary or energetically costly responses is critical to controlling infection without collateral damage. These diverse, dynamic responses are reflected in the transcriptome, making them an attractive substrate for computational classification.

A clear example of this functional specialization is seen in the differential response to DNA versus RNA viruses. For instance, infection with a DNA virus such as mpox (monkeypox) activates cytosolic DNA sensors like cGAS, DNA-PK, and IFI16, which initiate downstream signaling to induce interferon-β and other antiviral genes^13,14^. In contrast, RNA viruses such as SARS-CoV-2 or Influenza A are recognized primarily through double-stranded RNA intermediates by RIG-I and MDA5^52^. These sensors engage the mitochondrial adapter MAVS to trigger antiviral transcriptional programs^52^.

Transcriptomic studies have shown that even among related viral strains, host responses can differ markedly. For instance, in the 2022 outbreak of mpox (Clade IIb), several genes were significantly upregulated (e.g., SLC2A3, ATP2B1, VEGFA) or downregulated (MAP3K8, IL1A, SGK1), reflecting a distinct host signature^53^. Similarly, SARS-CoV-2 variants induce variant-specific responses: the Delta variant was associated with upregulation of EGR1 and IFIT, while Alpha triggered genes like SYVN1, CH25H, VIPR1, and others^54^.

In this study, we show that host transcriptomic data—whether represented as normalized gene expression values or alignment-free k-mer frequency spectra—contain sufficient information to classify the infecting virus using machine learning. For viruses like Influenza A, which contribute substantial amounts of viral RNA to the transcriptome and elicit strong transcriptional changes, both hierarchical clustering and Random forest classification achieved high performance. In contrast, even for viruses with more subtle effects on host gene expression and lower transcriptomic viral burden, Random forest classifiers were still able to differentiate viral genotypes with high accuracy. This likely reflects the ability of Random forests to capture non-linear patterns in complex, high-dimensional data^55^. Where hierarchical clustering failed to separate infected from uninfected samples—especially in cases where the host response was mild or variable—Random forest models consistently identified the correct viral infection. This suggests that while traditional clustering methods detect global patterns of perturbation, machine learning models can exploit finer-grained, distributed signals that are not apparent through unsupervised methods.

This study has several important limitations. Our analyses are based on infections in two immortalized human cell lines (Huh7 and Calu-3) under controlled laboratory conditions, which may not capture the complexity of in vivo or clinical infections. Host responses in primary cells, across tissues, or in whole organisms can differ in both timing and magnitude, potentially affecting classification performance. The datasets are further constrained to mid-stage infection (12–24 hpi); earlier or later responses may be weaker, noisier, or biologically distinct. In addition, our work is limited to negative-sense RNA viruses, leaving open how well this approach generalizes to positive-sense RNA or DNA viruses with different replication dynamics. Finally, clinical samples often present additional challenges—mixed infections, inter-individual variability, and technical artifacts such as batch effects or sequencing depth—that may reduce robustness. Although our depletion and enrichment experiments begin to address the role of viral read abundance, the minimal data requirements and cross-platform reproducibility remain unresolved.

Several directions could strengthen the translational relevance of this framework. Extending benchmarking to additional cell types, primary cultures, and eventually clinical samples will be essential for testing whether host-only signatures generalize beyond controlled in vitro settings. Longitudinal sampling across broader post-infection windows could help determine the earliest time points at which responses become diagnostic and the duration for which they remain informative..

Despite these caveats, our findings provide a clear proof-of-concept. We show that host transcriptomes alone are sufficient to distinguish between infections with diverse RNA viruses, even when viral reads are scarce and when unsupervised clustering fails. This underscores that host responses are not merely generic markers of infection but encode virus-specific, nonlinear signatures that can be captured by machine learning. In practice, such genome-agnostic signals could support early diagnostic triage when primers are unreliable or viral genomes are unknown, complementing sequence-based approaches. These results align with a growing body of work on transcriptomic diagnostics in bacterial sepsis and respiratory infections, suggesting a broader paradigm in which host responses serve as sensitive biosensors of pathogen class. By demonstrating consistent classification across multiple viral families, we add to evidence that systems-level host responses can bridge key gaps between pathogen discovery, outbreak surveillance, and clinical diagnostics.

## DATA AVAILABILITY

The data used in this study were obtained from the Sequencing Read Archive (PRJNA1074963) and Gene Expression Omnibus (GSE49840). From PRJNA1074963, we created synthetic permuted, shuffled, enriched, and depleted derivative datasets.

## MATERIALS AND METHODS

### Data sources

We analyzed publicly available RNA-seq datasets from human hepatocellular carcinoma (Huh7) and lung adenocarcinoma (Calu-3) cell lines infected with a panel of negative-sense RNA viruses, including influenza A subtypes (H1N1, H5N1, H7N7, H7N9), Ebola virus (EBOV), Marburg virus (MARV), Nipah virus (NiV), Rift Valley fever virus (RVFV), respiratory syncytial virus (RSV), Sandfly fever Sicilian virus (SFSV), and Lassa virus (LASV). Metadata on infection time point (3, 6, 7, 12 or 24 hpi), replicate, and sequencing depth were extracted from SRA and GEO project accessions (PRJNA1074963, GSE49840). We used the first 5,000 reads from each .fastq file, because we saw that the nucleotide k-mer spectra converged around 1000-5000 reads (Suppl. Fig. S1).

### Feature generation

We derived two complementary feature sets from host transcriptomes: (i) normalized differential gene expression (TPM) values and (ii) alignment-free k-mer frequency spectra (k = 1–15). Viral reads were excluded or enriched as controls by re-mapping and sub-sampling.

### Classification and clustering

To evaluate separability of host responses, we applied hierarchical clustering (Euclidean distance, Ward linkage) and Random Forest classifiers. For Random Forests, we used 200 trees, class-balanced stratified 4-fold cross-validation (smallest class n = 4), and default scikit-learn parameters unless otherwise specified.

### Controls

We implemented strict controls, including label permutation, read shuffling, viral read depletion, and viral read enrichment, to confirm that observed signal reflected biologically meaningful host responses.

### Reproducibility

All analyses were performed in Python (v3.11) using pandas, scikit-learn, Biopython, and SciPy. Data accessions are provided above; analysis code will be publicly archived on Zenodo before publication. All of the code is hosted on the following GitHub repository: https://github.com/blatuscaspot/multivirus-classification

## Supplemental Material

## Acknowledgements

We acknowledge the input of Dr. John Parker at Cornell University in suggesting to use shuffled nucleotides. This research was enabled in part by support provided by Calcul Québec (calculquebec.ca) and the Digital Research Alliance of Canada (alliancecan.ca).

