## Supplemental Figures for "Machine Learning Enables Viral Genome-Agnostic Classification of RNA Virus Infections from Host Transcriptomes"

### Machine Learning Reveals Nonlinear Transcriptomic Responses to Negative-Sense RNA Virus Infection

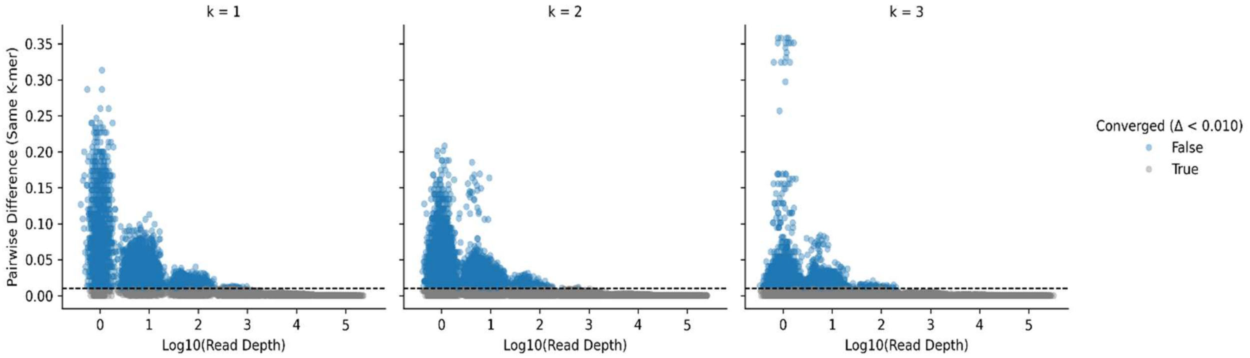


**Suppl. Fig. S1**: The kmer abundance profiles for low length kmers converges at about 1000 reads

**Supplementary Figure 1**: This figure shows how k-mer abundance estimates converge across sequencing replicates as read depth increases. Each point represents the pairwise difference in abundance for the same k-mer between two replicates, plotted against the log10 of the read depth. Differences below a threshold ($\Delta < 0.01$, dashed line) are classified as "converged" (grey), while larger differences are shown in blue. Across k-mer sizes, most estimates converge rapidly with increasing sequencing depth, with variance shrinking as read depth increases. This indicates that k-mer profiles stabilize and become reproducible across replicates at moderate sequencing depths.
